# Chronosequence and direct observation approaches reveal complementary community dynamics in a novel ecosystem

**DOI:** 10.1101/453217

**Authors:** Andrew Kulmatiski, Karen H. Beard

**Author notes:** (corresponding author,), http://orcid.org/0000-0001-9977-550.

## Abstract

Non-native, early-successional plants have been observed to maintain dominance for decades, particularly in semi-arid systems. Here, two approaches were used to detect potentially slow successional patterns in an invaded semi-arid system: chronosequence and direct observation. Plant communities in 25 shrub-steppe sites that represented a 50-year chronosequence of agricultural abandonment were monitored for 13 years. Each site contained a field abandoned from agriculture (ex-arable) and an adjacent never-tilled field. Ex-arable fields were dominated by short-lived, non-native plants. These ‘weedy’ communities had lower species richness, diversity and ground cover, and greater annual and forb cover than communities in never-tilled fields. Never-tilled fields were dominated by long-lived native plants. Across the chronosquence, plant community composition remained unchanged in both ex-arable and never-tilled fields. In contrast, 13 years of direct observation detected directional changes in plant community composition in both field types. Despite changes in community composition in both field types during direct observation, there was little evidence that native plants were invading ex-arable fields or that non-native plants were invading never-tilled fields. The more-controlled, direct observation approach was more sensitive to changes in community composition, but the chronosequence approach suggested that these changes are unlikely to manifest over longer time periods, at least in part because of disturbances in the system. Results highlight the long-term consequences of soil disturbance and the difficulty of restoring native perennials in disturbed semi-arid systems.

*Nomenclature*: Hitchcock and Cronquist (1973) for plants.

## Introduction

Over the past century, early-successional, non-native plants have invaded disturbed areas around the world (Richardson 2011; Jauni et al. 2015). Traditional ecological theory suggests that without continued disturbances late-successional species will gradually replace these invaders (Elton 1958; MacDougall & Turkington 2005; Myers & Harms 2009). Yet, in some, particularly semi-arid areas, early-successional, non-native species persist for decades or longer (Stylinski & Allen 1999; Courtois et al. 2004; Tognetti et al. 2010; Kachergis et al. 2014; Gill et al. 2018). Distinguishing whether early-successional, non-natives follow successional patterns or if they create alternate-state plant communities has important theoretical and management implications (Briske et al. 2005; Cramer et al. 2008; Murcia et al. 2014; Alsted et al. 2016). Where native perennial species re-establish in years or decades, management efforts may delay restoration (Rinella et al. 2009). Where alternate-state communities develop or where succession occurs at very slow rates (*i.e.,* centuries), intensive management approaches are likely needed (Cramer et al. 2008).

Assessments of long-term patterns of plant community composition are important for understanding how plant communities respond to disturbance and determining appropriate management approaches (Strayer 2012; Flory & D’Antonio 2015). The data needed to assess long-term community dynamics in semi-arid systems, however, is often lacking (Tognetti et al. 2010; Yelenik & D'Antonio 2013; Kachergis et al. 2014; Morris et al. 2014). Space-for-time substitutions (or chronosequences) can be used to infer species replacements over long periods. Chronosequence data, however, are susceptible to hidden temporal or spatial variations in factors such as climate, grazing, priority effects, and soil type (Foster & Tilman 2000; Bonet & Pausas 2004; Walker et al. 2010; Gill et al. 2018). Direct long-term observations of plant community dynamics can control for these extrinsic factors and provide a better test of community resistance and resilience to changes in species composition but are more difficult to collect (Blossey 1999).

Kulmatiski (2006) used a chronosequence approach to describe plant community dynamics in a sage-steppe ecosystem in Washington, USA. Results suggested that non-native and native plants established alternate-state communities in ex-arable fields and never-tilled fields, respectively. It was suggested that disturbance associated with agriculture allowed early-successional, non-native plants to establish, but that once established, these species changed soil conditions in ways that allowed their own persistence (*i.e*., positive plant-soil feedbacks; Kulmatiski 2006). To provide a more controlled test of how vegetation dynamics change over time, in the present study the same fields in that chronosequence were monitored for an additional 10 years to produce 13-year dataset of direct observation (Fukami & Wardle 2005). Direct observation allows an assessment of how the community in each individual field changes over time. Direct observation also allows an assessment of the effects of several extrinsic factors that were not possible to address during the previous study. During the course of this study, roughly half of the ex-arable fields were managed (*i.e*., tilled, herbicided and seeded) to increase native plant growth, most of the fields were burned in a wildfire in the penultimate year of surveying, and across all sites a biocontrol agent nearly eliminated the dominant non-native plant, *Centaurea diffusa* Lam.

We had three objectives in this study. First, we describe community differences in ex-arable and never-tilled fields using 13 years of direction observation data. Second, we use direct observation and chronosequence approaches to determine how the communities in the two fields are changing over time, and if they are changing, at what rate. Differences in the results between the two approaches are discussed. Third, we determine whether management and natural treatments change the trajectories of these communities.

## Methods

### Study area

Research was conducted in the Methow Valley, Washington (WA), USA (48° 37’ N, 120° 10’ W). Precipitation is seasonal with 250 of 360 mm of annual precipitation falling mostly as snow between October and March (http://www.ncdc.noaa.gov). The growing season begins with snowmelt in April and continues until snowfall in November though most native grasses and forbs become dormant by July. Native shrub-steppe communities dominated by *Purshia tridentata, Pseudoroegneria spicata*, *Lupinus sericeus*, *Artemesia tridentata*, and *Balsamorhizae sagittata* occupy most of the hilly landscape, whereas valley bottoms and benches are largely used for agriculture and, once abandoned, are occupied by various non-native species including *Centaurea diffusa* Lam*, Medicago sativa, Bromus tectorum, Poa bulbosa,* and *Cardaria draba*. Unless otherwise noted species naming follows that of Hitchcock and Cronquist (1973).

Aerial photographs were used to identify 25 study sites with fields that had been abandoned from low-input, dryland agriculture (*Triticum aestivum* and *Medicago sativa*) between 1950 and 1999 and that had adjacent undisturbed fields with similar slope, aspect and soils (S1 Table). Henceforth, these field types are referred to as ‘ex-arable’ and ‘never-tilled’, respectively. At least 200 m but not more than 25 km separated the 25 sites (elevation range: 630 to 1000 m). All sites were located on the Newbon-Concunully association (coarse-loamy, mixed mesic Typic Haploxerolls; Lenfesty 1980).

### Vegetation sampling

From 5 to 25 June from 2002 to 2015, plant cover by species was determined in 1 m x 1 m quadrats in each of 25 sites. Each of the 25 sites contained an ex-arable field and a paired never-tilled field. In each field, two transects were established. These transects were parallel to and 5 m or 50 m from the historical tillage boundary. Depending on field length, 15 quadrats were placed at an interval of one every 5 to 10 m in each transect. This sampling design (15 quadrats x 2 transect distances x 2 field types x 25 sites) produced 1,500 quadrats per year. Data was not available for the 2010 season, so 13 years of data are reported. The total dataset contains species compositions for 18,692 quadrats.

Plant cover was measured as percent ground cover using visual estimation by the same observer over the 13 years to the nearest 1% of cover. A 5 x 5 grid in the quadrat helped guide estimation. Due to the number of plots (1,500 each year) and the request of land managers, transects and plots were not permanently marked so that while fields and transects were resampled each year, specific plots were not resampled each year. Visual estimates of percent cover were well correlated with point-intercept-derived estimates of plant cover by species (i.e., R^2^ = 0.95 to 0.98; Kulmatiski 2006).

### Management and wildfire history

In 2003 and 2004, the biocontrol agent *Larinus minutus* (Gyllenahal) was released to control the dominant non-native, *Centaurea diffusa* Lam. Between 2005 and 2013, 12 of the ex-arable fields were managed to increase native plant abundance (Appendix Table 1). Management included broad-spectrum herbicide application in the spring followed by two to three passes with a disk harrow at two to four different times over two growing seasons (*e.g*., spring and fall) prior to a native plant seeding, typically in the fall. In the penultimate year of the study (August 2014), most (15 of the 25) of the sites burned in a wildfire (Appendix Table 1).

### Statistical analyses

To address our first objective, to test for differences between the two field types, we used non-metric multidimensional scaling analyses (NMS). NMS analyses were conducted on functional group (native annual, non-native annual, native forb, non-native forb, native grass, non-native grass, native perennial, non-native perennial, native shrub) and species matrices using Bray-Curtis dissimilarity matrices. Analysis of similarities (ANOSIM) was used to test for differences in NMS scores between treatments (*i.e*., ex-arable or never-tilled field; Clarke 1993; Sturrock & Rocha 2000).

To further address the first objective, we provide a more detailed analysis of differences in community composition between ex-arable fields and never-tilled fields, and between distance transects (5 and 50 m), using generalized linear mixed models (GLIMM) with a two-way factorial (field type by distance) in a split-plot design with whole plots (fields) in blocks (sites) and the following response variables: native, non-native, grass, forb, annual and perennial.

To address the second objective, to assess community composition change over time using both direct observation and chronosequence data, linear trend in the NMS1 scores over years-since-abandonment and comparison of trends were assessed using a linear mixed model. In addition to NMS1 scores, this analysis was done on functional groups and species (five dominant in each field type). Fixed effects factors were (1) field type (ex-arable or never-tilled); (2) years-since-abandonment; (3) mean number of years-since-abandonment; (4) interaction of field type with years-since-abandonment, which allowed the estimate of the within-field slope of the regression of NMS1 scores on number of years-since-abandonment to differ for ex-arable and never-tilled fields; and (5) interaction of field type with mean number of years-since-abandonment, which allowed the estimate of the between-field slope of the regression to differ for ex-arable and never-tilled fields (van de Pol and Wright 2009). Random effects factors were study site and field-within-study-site, which allowed for random intercepts for fields in the regressions of NMS1 scores on the number of years-since-abandonment. Covariance among the 13 annual repeated measures on a field was modeled using a first-order autoregressive structure. We also fitted a model that allowed for random slopes among field regressions; there was no statistical support for random slopes so we present results for the model with only random intercepts. Number of years-since-abandonment was within-field centered prior to analysis. For simplicity, only data from 50 m transects were analyzed.

To test for management effects in ex-arable fields, the third objective, regressions of NMS scores were conducted for managed and unmanaged fields separately. To test for biocontrol effects on the target non-native plant, *C. diffusa*, a one-way GLIMM with year as the factor and field as the random variable was used to test for differences in cover in each community. To test for the effects of wildfire, differences in NMS values in 2015 were compared to average NMS values from 2002 to 2014, using a t-test for each field type separately.

All multivariate analyses were performed in R using the LabDSV library (Roberts 2016; Team 2004; Appendix). Regressions and GLIMMs were conducted using the Reg and GLIMMIX procedures in SAS v.9.4 TS1M4 for Windows (SAS Institute, Cary, North Carolina, USA; Appendix).

## Results

### Community composition in ex-arable and never-tilled fields

NMS of the direct observation data revealed differences in functional group and species composition between ex-arable fields and never-tilled fields (ANOSIM statistic = 0.486, 0.444, *P* < 0.001, *P* < 0.001 for functional group and species matrices, respectively, Fig 1). NMS axis 1 scores distinguished differences in plant community composition between ex-arable fields and never-tilled fields, while NMS axis 2 scores distinguished differences in plant community composition among fields and the 13 years of observation (Fig 1).

**Fig. 1.**
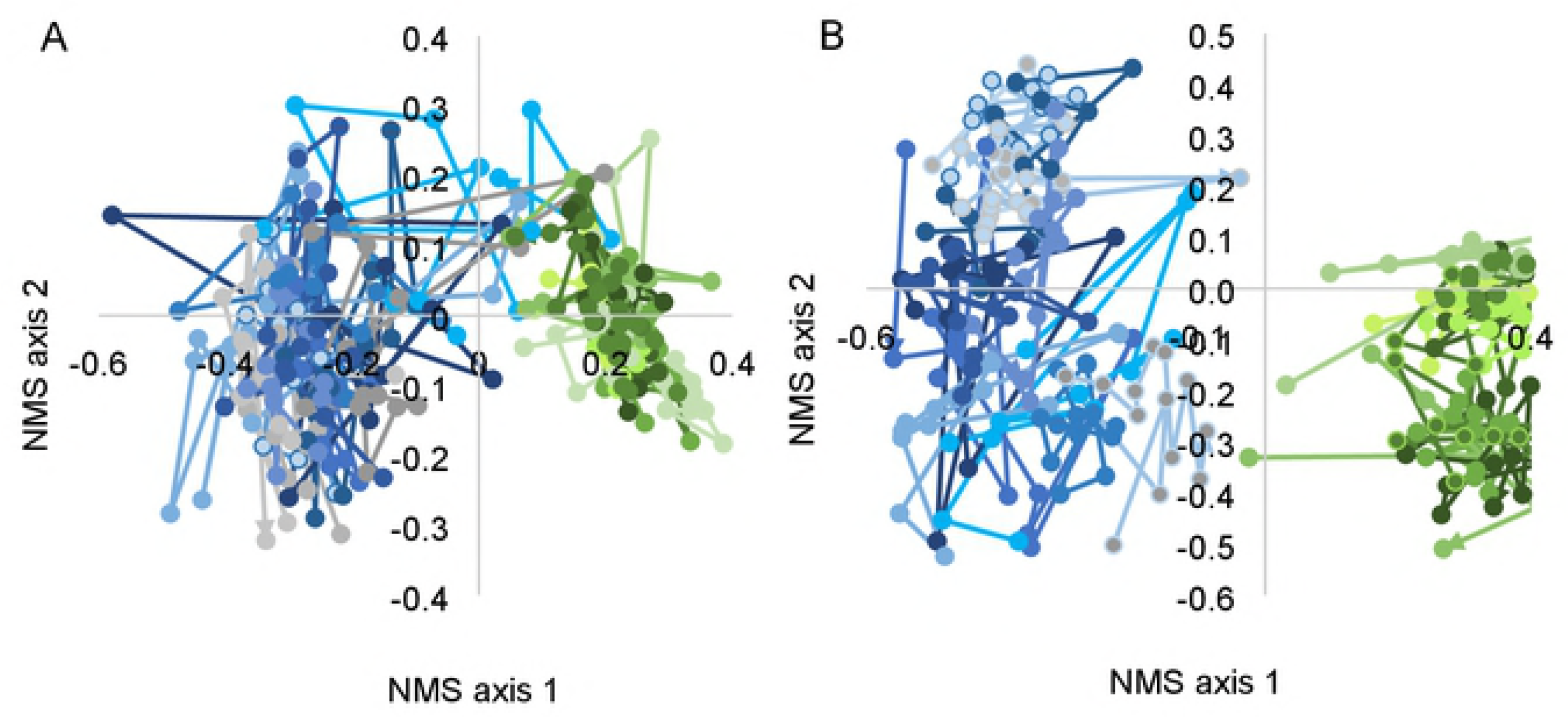
No nmetric multidimensional scaling (NMS) graph by (A) functional group and (B) species ofplant community composition in adjacent ex-arable fields (blue colors) and never-tilled fields (green colors) over 13 years. Each point represents the mean composition of vegetation across a transect located SO m from a tillage boundary. Lines connect values from a field over 13 years of direct observation. For clarity, data from 12 of25 randomly selected sites is shown.

Differences in plant functional group composition between ex-arable and never-tilled fields helped explain community-level differences indicated by NMS (Fig 2; Table 1). Non-natives, annuals and forbs were more abundant in ex-arable than never-tilled fields (Fig 2; Table 1). There was no difference in grass abundance between ex-arable and never-tilled fields.

**Fig. 2.**
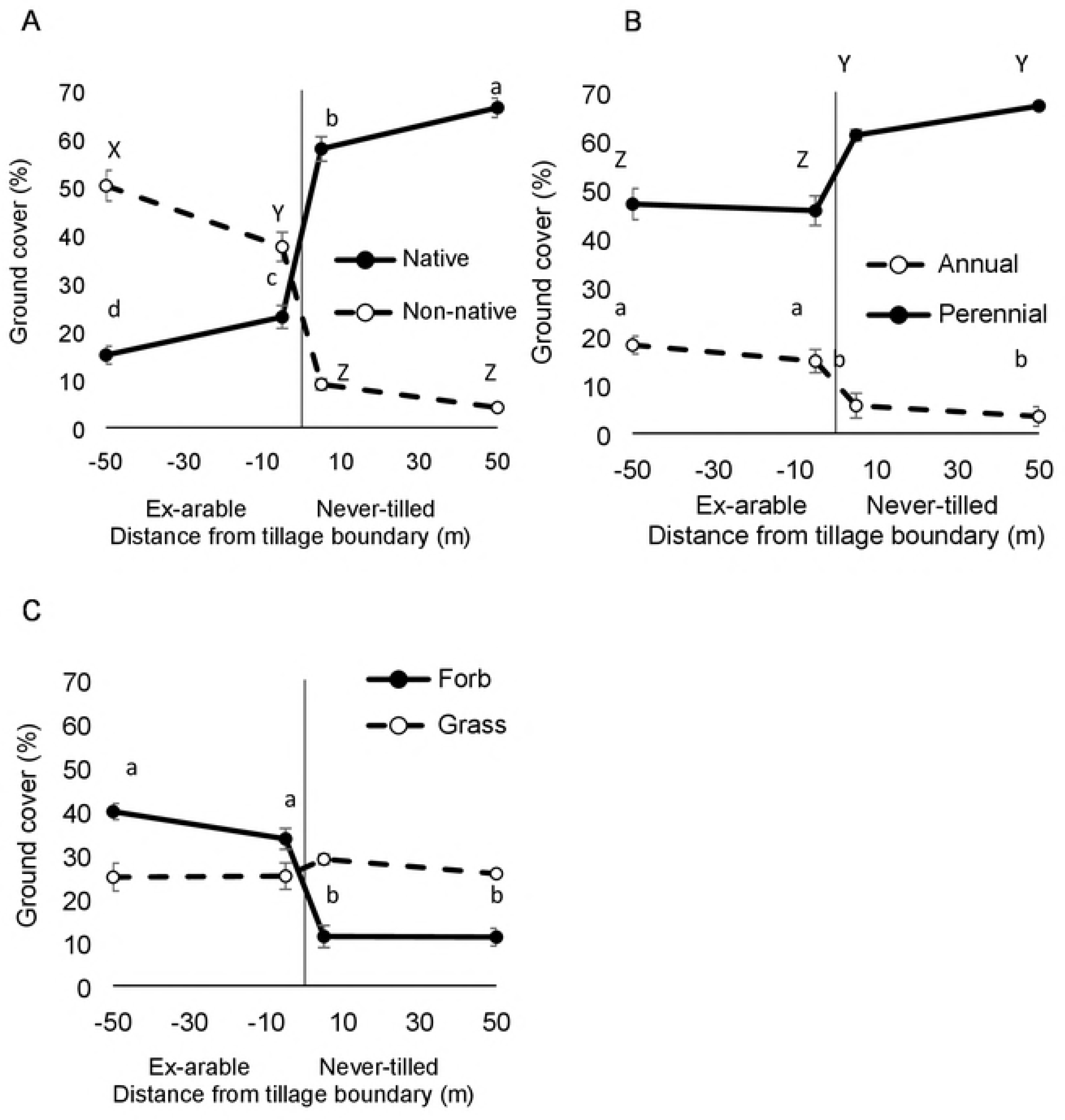
Percent ground cover of (A)native and non-native plants, (B)annual and perenialplants, and (C)forbs and grasses across tillage boundaries. Negative x-axis values are in ex-arable fields. Positive x-axis values are in never-tilled fields. Values represent the mean and standard error associated with the 25 sampled fields (values from replicate plots and years were averaged prior to calculations). Different letters are different at the α = 0.05 level.

**Table 1.** Mean cover of dominant plant species and functional groups and several community traits in adjacent ex-arable and never-tilled fields, Methow Valley, Washington, USA. Cover values within fields (replicate plots) and among years were averaged prior to calculations so that values represent averages and standard errors associated with the 25 fields.

| Measure | Ex-arable<br>fields | Never-tilled<br>fields |
| --- | --- | --- |
| Species richness | 5.4 ± 0.3 b | 6.6 ± 0.3 a |
| Ground Cover (%) | 59.2 ± 1.2 b | 68.6 ± 1.6 a |
| Native: | 19.2 ± 1.8 b | 62.4 ± 2.1 a |
| *Pseudoroegneria spicata* | 2.8 ± 1.2 b | 18.9 ± 1.3 a |
| *Balsamorhiza sagittata* | 0.4 ± 0.1 b | 15.3 ± 1.9 a |
| *Purshia tridentata* | 0.1 ± 0.0 b | 4.8 ± 0.9 a |
| *Lupinus* spp. | 2.3 ± 0.5 b | 4.3 ± 0.5 a |
| *Artemisia tridentata* | 0.1 ± 0.0 b | 2.4 ± 0.8 a |
| Non-native: | 39.6 ± 2.1 a | 6.6 ± 0.8 b |
| *Medicago sativa* | 7.1 ± 1.7 a | 0.3 ± 0.1 b |
| *Poa bulbosa* | 5.9 ± 0.9 a | 1.7 ± 0.3 b |
| *Centaurea diffusa* | 5.1 ± 0.8 a | 0.2 ± 0.1 b |
| *Bromus tectorum* | 4.5 ± 0.7 a | 1.4 ± 0.3 b |
| *Cardaria draba* | 4.3 ± 0.1 a | 0.1 ± 0.1 b |
| Native annual | 4.4 ± 0.7 b | 2.0 ± 0.2 a |
| Native forb | 7.9 ± 1.0 a | 8.2 ± 0.6 a |
| Native grass | 9.8 ± 1.5 b | 24.0 ± 0.6 a |
| Native perennial | $14.7 \pm 1.7$ b | $60.4 \pm 2.1$ a |
| Native shrub | $1.2 \pm 0.2$ b | $30.2 \pm 2.7$ a |
| Non-native annual | $11.3 \pm 1.6$ a | $2.6 \pm 0.5$ b |
| Non-native forb | $23.8 \pm 2.2$ a | $3.1 \pm 0.4$ b |
| Non-native grass | $15.8 \pm 1.6$ a | $3.5 \pm 0.5$ b |
| Non-native perennial | $28.3 \pm 2.4$ a | $4.0 \pm 0.5$ b |
| Non-native shrub | $0.0 \pm 0.0$ a | $0.0 \pm 0.0$ a |
| Annual | $15.8 \pm 1.8$ a | $4.6 \pm 0.6$ b |
| Forb | $31.7 \pm 2.3$ a | $11.3 \pm 0.8$ b |
| Grass | $25.7 \pm 1.8$ a | $27.4 \pm 1.3$ a |
| Perennial | $42.9 \pm 1.8$ b | $64.4 \pm 1.9$ a |
| Shrub | $1.2 \pm 0.2$ b | $30.2 \pm 2.7$ a |

<sup>1</sup> Values represent the average from thirty quadrats and 13 sampling years in each plant community in 25 fields (n = 25). The SE represents the error associated with the 25 fields and not error associated with quadrats, transects, or years.
<sup>2</sup> Values in the same row followed by the same letter were not significantly different at the 0.05 level.

### Community composition across time

There was evidence for an effect of field type (F_1,22_ = 44.44, *P* < 0.001) and effect of time during direct observation on plant community composition (F_1,78_ = 4.88, *P* = 0.030; Fig 3). The effect of time during direct observation was significant for ex-arable fields (F_1,78_ = 2.21, *P* = 0.030, slope = 0.007), but not for never-tilled fields (F_1,78_ = 0.00, *P* = 1.00). There was no effect of time for the chronosequence data (*i.e*., between-plot slope) for ex-arable (*F*_1,22_ = −1.28, *P* = 0.213) or never-tilled fields (*F*_1,22_ = −0.65, *P* = 0.520; Fig 3).

**Fig. 3.**
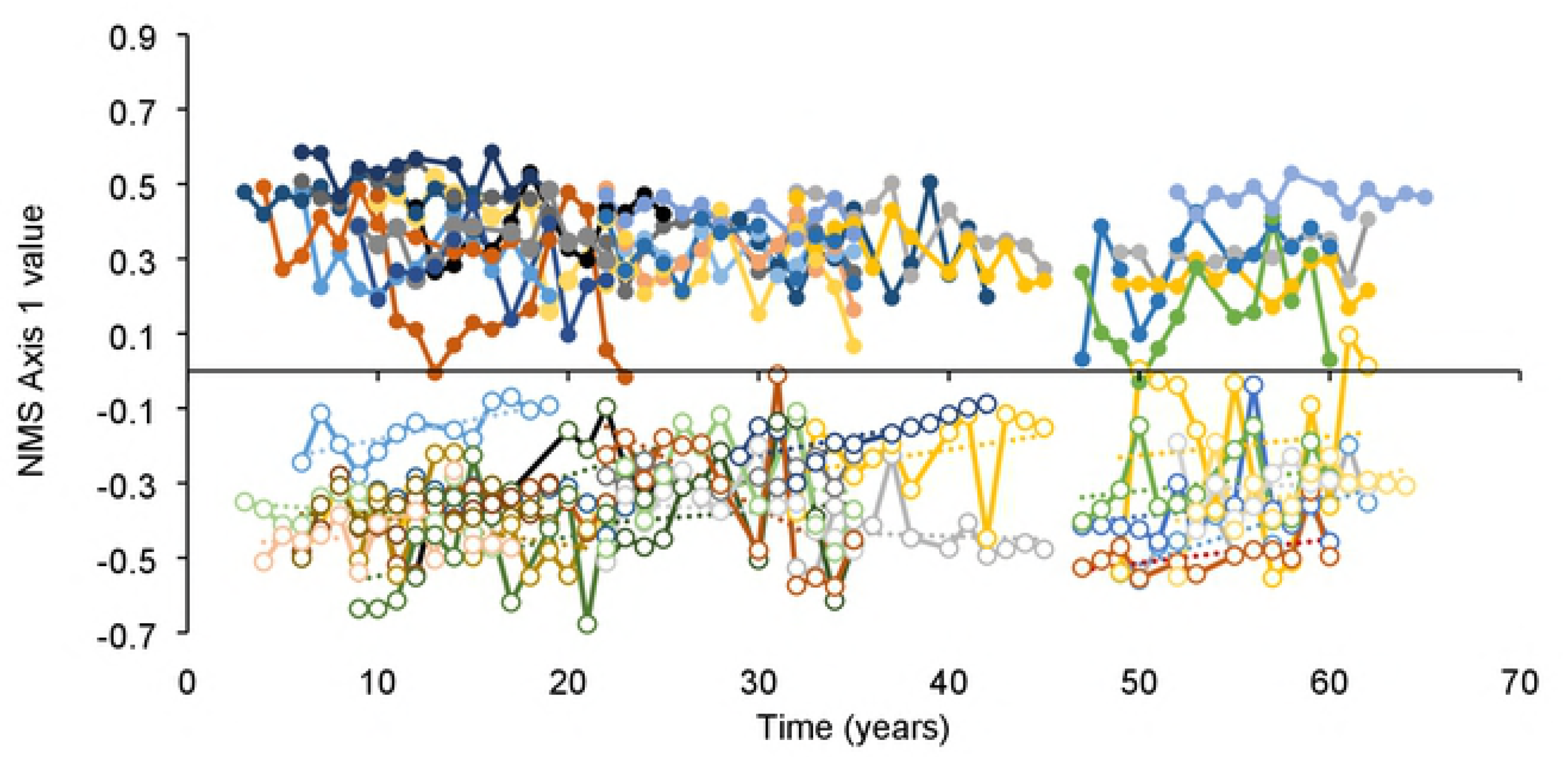
NMS n is 1 scores (species matrix) over time. Data from ex-arablefields shown with open symbols and data from never-tilled fields shown with filled symbols. Data from transects located 50 m from tillage boundaries. Positive y-axis values were associated with native, long lived perenial communites (Table 1; Fig. 1). NMS 1 values increased with time during 13 years of observation in ex-arable fields, but did not change with time in never-tilled communities or across the chronosequence in either community type.

Non-native plant abundance did not change across the chronosequence in ex-arable or never-tilled fields (F_1,23_ = 0.00, *P* = 0.974; S1 Fig), but non-native abundance did decrease in ex-arable fields during direct observation (F_1,48_ = 6.49, *P* = 0.014) at a rate of −0.84% cover per year (S1 Fig). Native plant abundance did not change across the chronosequence (F_1,46_ = 0.75, *P* = 0.391), but native abundance decreased in never-tilled fields during direct observation (F_1,46_ = 7.2, *P* = 0.010) at a rate of −0.67% cover per year (S1 Fig).

Among native plant species, *A. tridentata* cover did not change during direct observation (F_1,106_ = 0.02, *P* = 0.895; Appendix Fig 2) or the chronosequence (F_1,11_ = 4.07, *P* = 0.069; data not shown). *B. sagittata* cover did not change during direct observation (F_1,86_ = 1.54, *P* = 0.218) or the chronosequence (F_1,11_ = 0.67, *P* = 0.432). *L. sericeus* cover increased in ex-arable fields during direct observation (F_1,69_ = 3.28, *P* = 0.002, slope = 0.23% per year) but not during the chronosequence (F_1,22_ = 0.00, *P* = 0.99). *P.* spicata cover decreased in never-tilled fields during direct observation (F_1,108_ = 7.59, *P* = 0.001, slope = −0.35% per year) but increased across the chronosequence (F_1,21_ = 2.66, *P* = 0.015, slope = 0.174). *P.* tridentata cover decreased during direct observation in never-tilled fields (F_1,48_ = 19.21, *P* < 0.001, slope = −0.42% per year), but not across the chronosequence (F_1,46_ = 0.17, *P* = 0.678).

Among non-native plants, *B. tectorum* cover decreased during direct observation in ex-arable fields (F_1,98_ = 5.97, *P* = 0.016; slope = −0.10; S2 Fig) and the chronosequence (F_1,22_ = 4.30, *P* = 0.050; slope = −0.004% per year; data not shown). *C. diffusa* Lam. cover did not change during direct observation (F_1,124_ = 0.66, *P* = 0.509), but decreased across the chronosequence in ex-arable fields (F_2,30_ = 5.16, *P* = 0.012; slope = −0.13% per year). *C. draba* cover did not change during direct observation (F_1,48_ = 2.00, *P* = 0.164) or the chronosequence (F_1,23_ = 0.65, *P* = 0.43). *M. sativa* cover did not change across direct observation (F_1,48_ = 0.35, *P* = 0.558) or the chronosequence (F_1,23_ = 1.15, *P* = 0.295). *P. bulbosa* cover decreased during direct observation in both field types (F_1,48_ = 5.58, *P* = 0.022; slope = −0.277), but not across the chronosequence (F_1,46_ = 3.43, *P* = 0.070).

### Community composition responses to management, biocontrol and wildfire

When data from managed and unmanaged ex-arable fields were analyzed separately, communities in unmanaged fields became more similar to never-tilled fields while communities in managed fields did not show a directional change in composition [i.e., NMS axis 1 values increased with time in unmanaged fields (F_1,11_ = 18.38, *P* = 0.001), but not in managed fields (F_1,10_ = 2.07, *P* = 0.18; data not shown)]. In response to biocontrol treatment, cover of the target, *C. diffusa* Lam., decreased to near zero in 2003 and 2004 and increased to become a dominant species in 2014 and 2015 (S2 Fig). In ex-arable fields, NMS axis 1 scores did not differ before and after wildfire (T_23_ = 0.43, *P* = 0.67, Fig 4). However, in never-tilled fields, NMS axis 1 values were smaller after wildfire indicating that these fields became more like ex-arable fields (T_23_ = 3.90, *P* < 0.001, Fig 4).

**Fig. 4.**
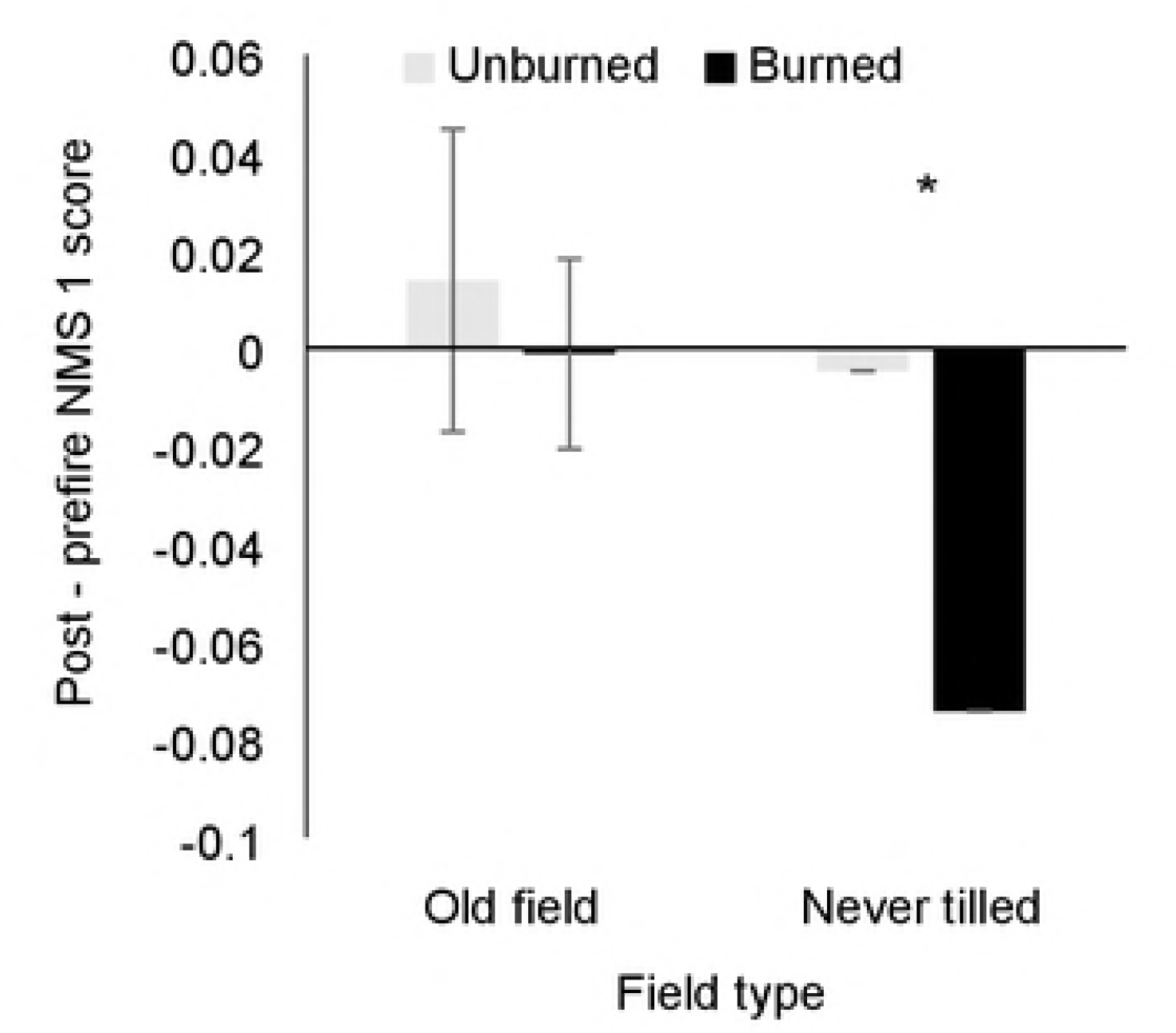
Change in NMS axis 1 values after a wildfire in 2014. Values calculated as the difference between 2015 scores and the average of 2002 to 2014 values. Positive NMS axis 1 scores were associated with weediness (Fig. 1). An asterisk indicates a difference between Unburned and Burned values at the α = 0.05 level.

## Discussion

Agricultural abandonment creates two distinct plant communities in the study system. Fields that have been disturbed by agriculture and abandoned are dominated by short-lived non-natives. Never-tilled fields are dominated by long-lived natives. Thirteen years of direct observation revealed directional changes in both field types. Non-native abundance decreased in ex-arable fields while native abundance decreased in never-tilled fields. These changes made the two communities more similar, but there was little evidence that the two community types would eventually converge because there was little evidence of species exchange between field types. Further, there was no evidence of directional changes in community composition across the chronosequence. In summary, the more-controlled, direct observation approach detected changes in community composition over 13 years, but the chronosequence approach suggests that these changes are unlikely to be realized over longer time periods.

Communities in ex-arable and never-tilled fields are distinct (Fig 1). In ex-arable fields, the five most abundant species are non-native, typically live a maximum of 1 to 10 years (Rumbaugh 1982) and demonstrate wide fluctuations in percent cover from year to year (Fig 1 and S2 Fig). These ‘weedy’ communities have fewer species, less perennial cover, and more annual and forb cover than communities in never-tilled fields. In contrast, the five most abundant species in never-tilled fields are native, often live more than 10 years (Rumbaugh 1982), and demonstrate more stable abundances over annual and decadal time scales.

While plant communities in ex-arable fields remained distinct from communities in never-tilled fields for 65 years, there was some evidence of directional change in community composition in the 13 years of direction observation. Multivariate analyses suggested that plant communities in ex-arable fields were becoming more similar to plant communities in never-tilled fields (i.e., NMS 1 values increased in ex-arable fields during direct observation; Fig 3). This was consistent with a decline in the abundance of *B. tectorum* and *P. bulbosa.* Both species are small-statured, winter-active, non-native grasses that are dominant in ex-arable fields but uncommon in never-tilled fields. The only evidence that native species were becoming more abundant in ex-arable fields over time was for the N-fixing *L. sericeus*; total native cover in ex-arable fields did not change during direct observation or the chronosequence. Thus, while two common non-natives decreased and one native species increased, there was little evidence that ex-arable communities would return to native community composition over 50 to 100 year timescales. In contrast to ex-arable fields, there was no directional change in whole-community composition in never-tilled fields during direct observation. At the functional group and species-levels, however, there were declines in native plant abundance reflecting a decline in the common shrub, *Purshia tridentata.* There was no evidence that non-native plants were invading never-tilled fields. Broadly, direct observation data appeared sensitive to detecting within-community changes in species composition, but chronosequence data suggested that communities will remain dissimilar over long (60+) time periods.

There are several reasons why direct observation data may have revealed different patterns in community composition than chronosequence data (Johnson and Miyanishi 2008). First, unlike chronosequence data, direct observation data are not confounded by space-for-time substitutions. For example, differences in community composition among fields can ‘mask’ within-community changes over time in chronosequence datasets (Foster & Tilman 2000; Bonet & Pausas 2004; Walker et al. 2010; Gill et al. 2018). Second, shorter-term, direct observation data are more likely to detect short-term linear patterns, even if longer-term patterns are non-linear. It is likely, for example, that weedy communities shift from annual to perennial dominance over five to ten year timescales, but that these communities then maintain dominance by perennial non-natives indefinitely. While the direct observation approach appeared more controlled and more sensitive to detecting community level changes, the inference it provides to community composition in the future is not as strong as from chronosequence data. It is possible, for example, that patterns observed in the direct observation dataset will be periodically ‘reset’ by disturbances from pocket gophers, drought, wildfire or human disturbance in ex-arable fields (Kyle et al. 2006; Gill et al. 2018). In any case, the patterns observed during direct observation are unlikely to be maintained over longer time periods because 1) these directional changes were not observed in the chronosequence and 2) plant cover was observed to decline in both field types during direct observation, but plant cover cannot be expected to decline indefinitely.

It is surprising that native, late-successional species did not recolonize ex-arable fields (Csecserits et al. 2001; Bonet & Pausas 2004; Meiners 2007). Ex-arable fields are typically less than 200 m wide allowing native propagule pressure and results were similar in transects that were 5 m and 50 m from tillage boundaries. Ex-arable fields had less ground cover and more variable populations, which should have provided establishment opportunities for many generations of plants (Richardson & Pyšek 2006; Gill et al. 2018). Species in ex-arable fields likely experienced 30 to 60 generations during the chronosequence (e.g., *C. draba* and *M. officinale*; Rumbaugh 1982; Dietz & Schweingruber 2002; Lauenroth and Adler 2008; Chu et al. 2014). Even the longest-lived native shrubs in the study system, *A. tridentata* and *P. tridentata*, have mean ages less than 25 years, and realize recruitment events every 2 to 3 years (Krannitz & Hicks 2000; Barry et al. 2001; Maier et al. 2001; Ziegenhagen & Miller 2009; Schlaepfer et al. 2014). Thus, establishment, recruitment and mortality were expected in both the direct observation and chronosequence data in both communities (Dalgleish et al. 2011), yet these communities remained distinct for 65 years.

Results stand in contrast to many studies demonstrating succession over similar time-scales (Foster & Tilman 2000; Csecserits et al. 2001; Bonet & Pausas 2004; Keeley et al. 2006; Meiners 2007; Li et al. 2016). It is not clear why succession would be rapid in some systems and not in others (Didham et al. 2005), though arid and semi-arid systems seem to be more likely to show very slow to no change (Stylinski & Allen 1999; Jackson & Bartolome 2002; Kulmatiski 2006; Kachergis et al. 2014) relative to more mesic systems (Rejmanek 1996; Bonet & Pausas 2004; MacDougall & Turkington 2005; Sojneková & Chytrý 2015). It is possible that slow succession or alternate-state communities are more likely in more stressful environments with greater facilitation (Flory & Bauer 2014; He & Bertness 2014; Michalet et al. 2014).

Direct observation data provided some insight into the processes through which short-lived non-natives may maintain dominance. It is particularly notable that the dominant species in ex-arable fields were resilient from changes in abundance. Perhaps the most dramatic example was that *C. diffusa* Lam. abundance declined to almost zero following the introduction of a biocontrol agent in 2003, then again become the dominant non-native species in 2014 and 2015. The temporary loss of *C. diffusa* Lam. was notable, but it was not the only source of variation in species abundances in ex-arable fields; most of the dominant species in ex-arable fields demonstrated large year-to-year changes in abundance. In addition to being resilient from changes in abundance, ex-arable communities were resistant to changes caused by management and wildfire. Active and intensive management failed to shift ex-arable communities toward the composition of never-tilled fields and wildfire had no effect on ‘weediness’ in ex-arable fields (Fig 4). Together, these results illustrate that ex-arable plant communities were either resistant to or resilient from large disturbances (*i.e*., biocontrol addition, native plant restoration efforts or wildfire; Klinger et al. 2017; Monaco et al. 2017).

Previous research at this and other sites suggests that facilitation among non-natives is more likely than agricultural legacies in explaining non-native persistence in ex-arable fields. For example, soil biological and nutrient traits were better associated with plant origin than agricultural history (Kulmatiski et al. 2006; Kulmatiski & Beard 2008). Once established in disturbed soils, non-native plants have been found to 1) create unique soil microbial communities (Kulmatiski et al. 2006; Stark and Norton 2015), 2) create soils with low penetration resistance (Kyle et al. 2007; Kyle et al. 2008), and 3) change water-use patterns in ways that inhibit the germination and establishment of native plants (Warren et al. 2015). In short, non-natives appear to create an environment that facilitates the growth of short-lived species and delays succession.

## Acknowledgements

This research was supported by the Utah Agricultural Experiment Station, Utah State University. Research was also supported by the National Science Foundation Award #1354129. Land use was approved by the Washington Department of Fish and Wildlife. We thank WDFW land managers, Jim Mountjoy, Kim Romain-Bondi, and Tom McCoy for their support with this research.

## Data accessibility

Upon acceptance, data will be made available at Utah State University Digital Commons (https://digitalcommons.usu.edu/all_datasets/) and by contacting the author.

